# A generic normalization method for proper quantification in untargeted proteomics screening

**DOI:** 10.1101/307504

**Authors:** Sandra Isabel Anjo, Isaura Simões, Pedro Castanheira, Mário Grãos, Bruno Manadas

## Abstract

The label-free quantitative mass spectrometry methods, in particular, the SWATH-MS approach, have gained popularity and became a powerful technique for comparison of large datasets. In the present work, it is introduced the use of recombinant proteins as internal standards for untargeted label-free methods. The proposed internal standard strategy reveals a similar intragroup normalization capacity when compared with the most common normalization methods, with the additional advantage of maintaining the overall proteome changes between groups (which are lost using other methods). Therefore, the proposed strategy is able to maintain a good performance even when large qualitative and quantitative differences in sample composition are observed, such as the ones induced by biological regulation (as observed in secretome and other biofluids’ analyses) or by enrichment approaches (such as immunopurifications). Moreover, this approach corresponds to a cost-effective alternative, easier to implement than the current stable-isotope labeling internal standards, therefore being an appealing strategy for large quantitative screening, as clinical cohorts for biomarker discovery.

## INTRODUCTION

Clinical assays and large screenings are highly informative studies that significantly contribute to expand scientific knowledge given their ability to combine information acquired from large cohorts of samples. The development of new label-free mass spectrometry methods able to analyze these large number of samples resulted in the establishment of different MS approaches, with the SWATH-MS emerging as a powerful technique for clinical applications and large screenings [1, 2]. However, in most of these studies samples are analyzed separately making these methods highly dependent on run-to-run reproducibility [3, 4]. Therefore, the use of internal standards (IS) - which are well stablished in targeted methods as one of the most important tools to obtain an accurate quantification [5] - corresponds to a valuable tool to correct for analytical variability and normalize data between experiments.

When only few molecules are analyzed, the most commonly used IS strategy is stable-isotope labeled (SIL) alternatives of the molecule of interest. This will induce a mass shift in the IS allowing the distinction between the IS and the analyte, while no major changes are induced in the remaining biochemical properties [5, 6]. Alternatively, strategies involving the labeling of entire proteomes have been applied for the quantification of a large number of proteins. In these methods, termed as global/whole proteome internal standard strategies, the labeled proteome is diluted into the samples and protein quantification is performed between the pair of unlabeled and labeled peptides [7, 8]. However, for untargeted proteomics a large variety of peptides can be generated and analyzed and, therefore, the use of an isotopically labeled standard may not properly represent sample heterogeneity. Moreover, the global proteome internal standard leads to an increase in sample complexity, resulting in a reduction in the number of protein identified in a single analysis. Additionally, the preparation of isotope-labeled compounds is time-consuming and expensive [9], precluding their use in routine analysis or in high throughput screenings.

Less common alternatives, include the use of analogous proteins [10] (targeted semi-quantitative applications) or the use of “housekeeping”-like proteins (pseudo internal standards) [9]. However, considering the multitude of proteomics samples, the use of a universal “housekeeping” proteins may be very challenging. Based on these limitations, the use of IS is residual in large untargeted label-free proteomics screenings, being substituted by the application of normalization methods, generally used in genomics and transcriptomics microarray analysis [11]. These normalization methods correspond to mathematical algorithms that are dependent on properties/features of the dataset. Thus, different methods may be applied to different types of screening [11, 12] making it impossible to identify a standard method.

In this work, it is proposed the use of non-labeled recombinant proteins as internal standard (IS) for untargeted methods as a cost-effective and transversal alternative to monitor technical variation caused by sample processing. Moreover, it is also addressed the best proteomics strategy to apply in samples where large differences are expected in protein content. The use of proteins instead of peptides will allow to monitor sample processing prior to protein digestion, and to obtain a large variety of peptides with the main properties of the peptides usually produced by the specific enzyme used in the assay. Since enzyme-specific proteolytic peptides prone to be used for quantification will be produced in all the experiments, the proposed IS might theoretically be used independently of the enzyme. The generated peptides will have different masses, as well as different retention times, being better estimators of the complex mixtures of peptides analyzed in untargeted proteomics studies and, thereby, allowing both samples normalization and adjustment of the retention time required in label free approaches.

A mixture of two recombinant proteins, the green fluorescent protein (GFP) and the maltose-binding protein (MBP), was evaluated as internal standards in a label-free proteomics screening. To establish their use as IS for untargeted screenings some parameters must be satisfied, namely: (1) the proteins and the tryptic peptides used must have no sequence similarity with any other protein from the sample; (2) the monitored peptides must be reproducibly quantified, and (3) the use of the IS must be able to accommodate for sample variability improving the precision of the results while leading to an accurate quantification. Those criteria were assessed using different strategies starting by a proteomics characterization of the internal standard mixture, evaluation of the impact of sample processing in the quantification of the IS, and finally comparing its normalization ability against a well-established normalization method.

## RESULTS

### Proteomics characterization of the internal standard mixture

The IS mixture proposed in this work was constituted by the recombinant proteins GFP (originally isolated from the jellyfish *Aequorea victoria* [13]) and MBP (part of the maltose/maltodextrin system of gram-negative bacteria [14]), both widely used as tools of genetic engineering: GFP is mainly used as a reporter for *in vivo* study of protein translocation [13], and MBP as a solubility enhancement tag for recombinant protein production, as well as its maltose binding activity used in protein affinity purification [15]. Sequence alignment against the entire reviewed UniProt database (Supplementary Table 1) further confirmed that both GFP and MBP only share sequence similarity to a restricted number of proteins from similar species and not to the proteome of the mammalian organisms most commonly used in proteomics screenings (human, mouse and rat species). Thus, these recombinant proteins may be easily distinguished from the majority of proteomics samples (at least of mammalian origin), confirming their suitability as internal standard candidates.

Additionally, mass spectrometry characterization of the IS mixture (Supplementary Table 2 and Supplementary Figures 1–2) revealed that none of the generated peptides are shared with another protein from the UniProt reviewed database (without species specific restrictions), indicating that the quantification of the targeted proteins will not be affected by the presence of the IS peptides, neither the quantification of the IS peptides will be prejudiced by the matrix (Supplementary Figure 3).

### Reproducibility analysis of internal standard quantification

A good IS has to be consistently quantified to be able to reflect the analytical variability induced by sample processing if significant variations occur. Therefore, it must have a reproducible quantitative performance, not affected by the main steps of the analysis, namely: (i) mass spectrometry acquisition; (ii) sample cleanup prior to mass spectrometry; and more importantly (iii) protein digestion. Additionally, in untargeted methods, the sample composition - particularly differences in sample complexity [9] - may lead to an additional source of variability to which the IS might be unresponsive.

Consequently, the impact of all the referred steps on the quantification of the proposed IS was assessed by a set of three specific experiments (Figure 1a). The impact of sample complexity was addressed in allexperiments by analyzing the IS alone, or in the presence of different amounts of a rat protein extract (“-/+ extract” conditions). Except for the experiment where the impact of protein digestion was evaluated, a single digestion of the IS solution and the protein extract was performed, and samples were combined at the peptide level immediately before the step being evaluated. In the LC-MS/MS evaluation experiment, the variability at the acquisition level was assessed by multiple injections of (i) the IS peptides solution alone, or (ii) the mixture of IS peptides with peptides from the protein extract prepared with samples previously cleaned. In the case of sample cleanup by solid phase extraction (SPE) using micropipette tips with C18 stationary phase, a single mixture was prepared and then divided into a total of four technical replicates to be subjected to parallel sample cleanup. Technical replicates were used to reduce biological variability that could mask the desired result.

**Figure 1.**
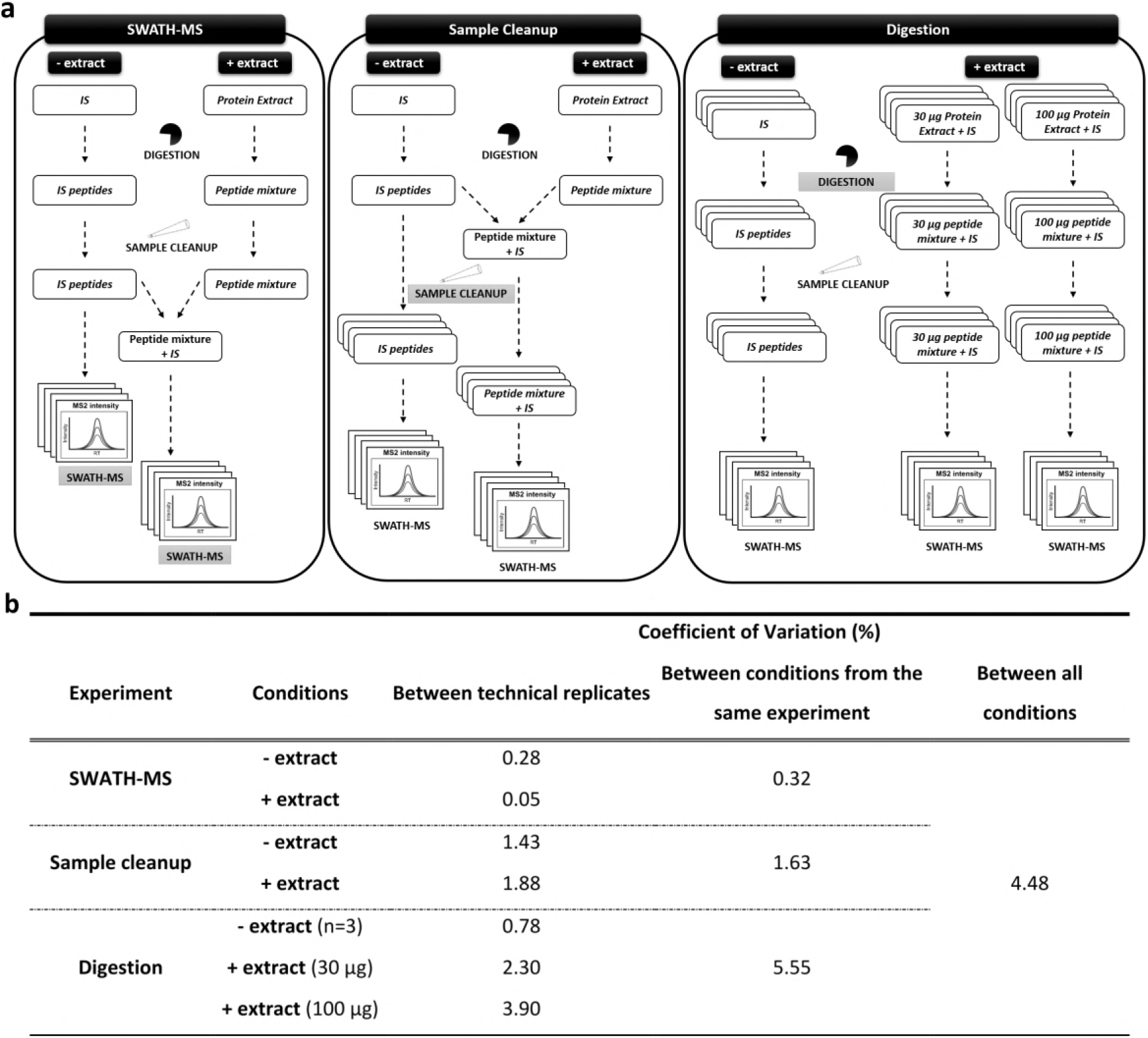
Impact of sample processing in the quantification of the proposed internal standard (IS) mixture. **(a)** Experimental workflow used to evaluate the consistency of the IS quantification. The impact of the three major steps in proteomics analysis (LS-MS/MS analysis, sample cleanup and protein digestion) on IS quantification was assessed by three independent experiments (separated by the rounded squares with the evaluated step highlighted). The impact of samples complexity was also evaluated by comparing the IS alone with a mixture of the IS in a rat protein extract (“-/+ extract” conditions, respectively), and by the impact of two different amounts of protein extract (30 *versus* 100 μg) during protein digestion. To avoid biological variability, technical replicates were used, in a total of four replicates per condition. Except for the experiment to evaluate protein digestion, a single digestion of the IS or the protein extract was performed, and their mixtures were done at the peptide level immediately before the step being evaluated. Quantitative analysis was performed by SWATH-MS/MS, and the internal standard quantification was evaluated by determination of the coefficient of variation (CV) **(b)**. The CV analysis between the different conditions per experiment indicates the impact of sample complexity in the quantification of the IS. Since the same amount of IS was added to all conditions, the final comparison between all the different conditions is an indication of the robustness of the IS quantification to different sample processing.

The quantification was performed by SWATH-MS/MS essentially as described in Anjo *et al.* [16], using a library composed by a combination of proteins identified from the rat extract with the two proteins used as internal standards (GFP and MBP). A total of 14 peptides from both recombinant proteins (GFP and MBP; Supplementary Table 3) were used to estimate the IS levels, which corresponds to the sum of the chromatographic peak areas of the fragments of the referred peptides [16]. It is expected that the IS may reflect some of the impact of technical variation. Therefore, to address only the impact induced in the measured values of the IS, and not the expected technical variation (such as difference in the sample or injection volume), the IS values were further normalized by the total intensity of the elements common to all samples [the internal standard and the iRT peptides (Biognosys) spiked prior to LC-MS/MS analysis].

The consistency of the quantification of the proposed IS mixture was determined by analyzing the coefficient of variation (CV) of the technical replicates (Figure 1b). Additionally, because the amount of IS was the same in all the conditions, it was also possible to evaluate the impact of sample complexity. In this case, the CVs were also determined between the “-/+ extract” conditions from the same experiment, and ultimately between all the experimental conditions. The largest CV variations were observed with the digestion of large amounts of proteins (CV of 5.5%) but remain below the 10% CV expected for technical replicates. Moreover, a similar percentage of CV was obtained when all the conditions were considered, indicating that the quantification of the proposed IS was very robust, since the effect of sample processing and/or complexity was minimal.

In addition to the observed reproducibility of the IS quantification, the peptides used as IS correlate well with a common mass-to-charge (*m/z*) distribution of a representative number of quantified peptides (Supplementary Figure 4 and Supplementary Table 3), being valuable representatives of the typical sample’s heterogeneity. Finally, these peptides are also well distributed across the chromatographic gradient (see Supplementary Table 3), allowing both the normalization of the protein levels and the adjustment of the retention times (RT), a key issue in label-free analysis.

### Evaluation of the normalization capacity of the IS

The IS must be able to reflect and accommodate sample-processing errors, and thus allow proper protein quantification between replicates and between different experimental conditions. In order to evaluate that capacity, the normalization using the proposed IS was compared against a well-established normalization method, the total intensity (TI) [12] in two different experiments. In the first experiment (Figure 2a) it was evaluated (i) the impact of high-demanding sample processing (herein represented by protein precipitation with TCA) and (ii) the differences in sample complexity (induced by distinct amounts of total protein extract) on the quantification reproducibility of a representative number of proteins. Technical replicates of the protein extract were used, to reduce the significant impact of biological variability on data reproducibility, but also to ensure that the acquired data could be successfully normalized by the TI methods (Figure 2b), and thus allow to properly evaluate the normalization capacity of the proposed IS mixture.

**Figure 2.**
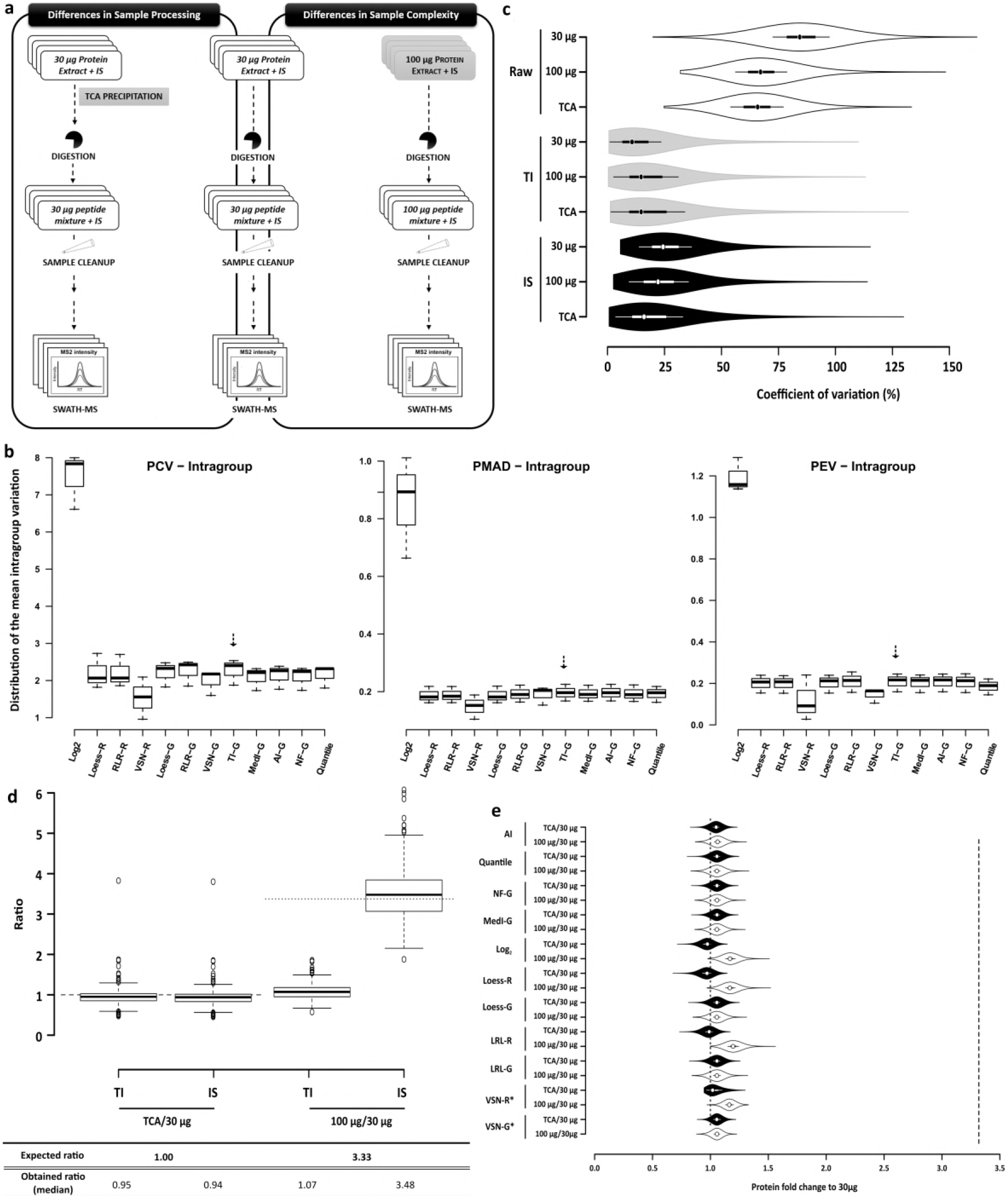
IS normalization reduces the intragroup variability while maintaining the intergroup variation. **(a)** Experimental workflow. The normalization capacity of the IS was tested by the analysis of the reproducibility of a representative number of proteins quantified in a protein extract. The impact of both sample processing and sample complexity was addressed (indicated by the boxes, with the correspondent evaluated parameter highlighted). To avoid biological variability, technical replicates were used, n=4. **(b)** Determination of the intragroup variation in the conditions presented in (a) using 12 normalization methods widely used in untargeted proteomics (see Online Methods for details). Arrows indicate the results for total intensity (TI). **(c)** Violin plots representing the distribution of the coefficient of variation with or without normalization (raw data; white polygons) to the TI (grey polygons) or the IS (black polygons). **(d)** Box plot representing the distribution of the protein fold change to 30 μg condition (used to assess the intergroup variation). The expected ratios for TCA/30 μg (dashed line) and 100 μg/30 μg (dotted line) and the obtained ratios are indicated in the bottom panel. **(e)** Violin plot representing the distribution of the protein fold change to 30 μg condition using the remaining 11 normalization methods from (b). * - method that leads to a lower intragroup variation according with the results from (b), which should be the best method to use in the dataset. Expected fold values for TCA/30 μg (black polygons; dotted line) should be 1 and 3.3 for 100 μg/30 μg ratio (white polygons; dashed line).

**Figure 3.**
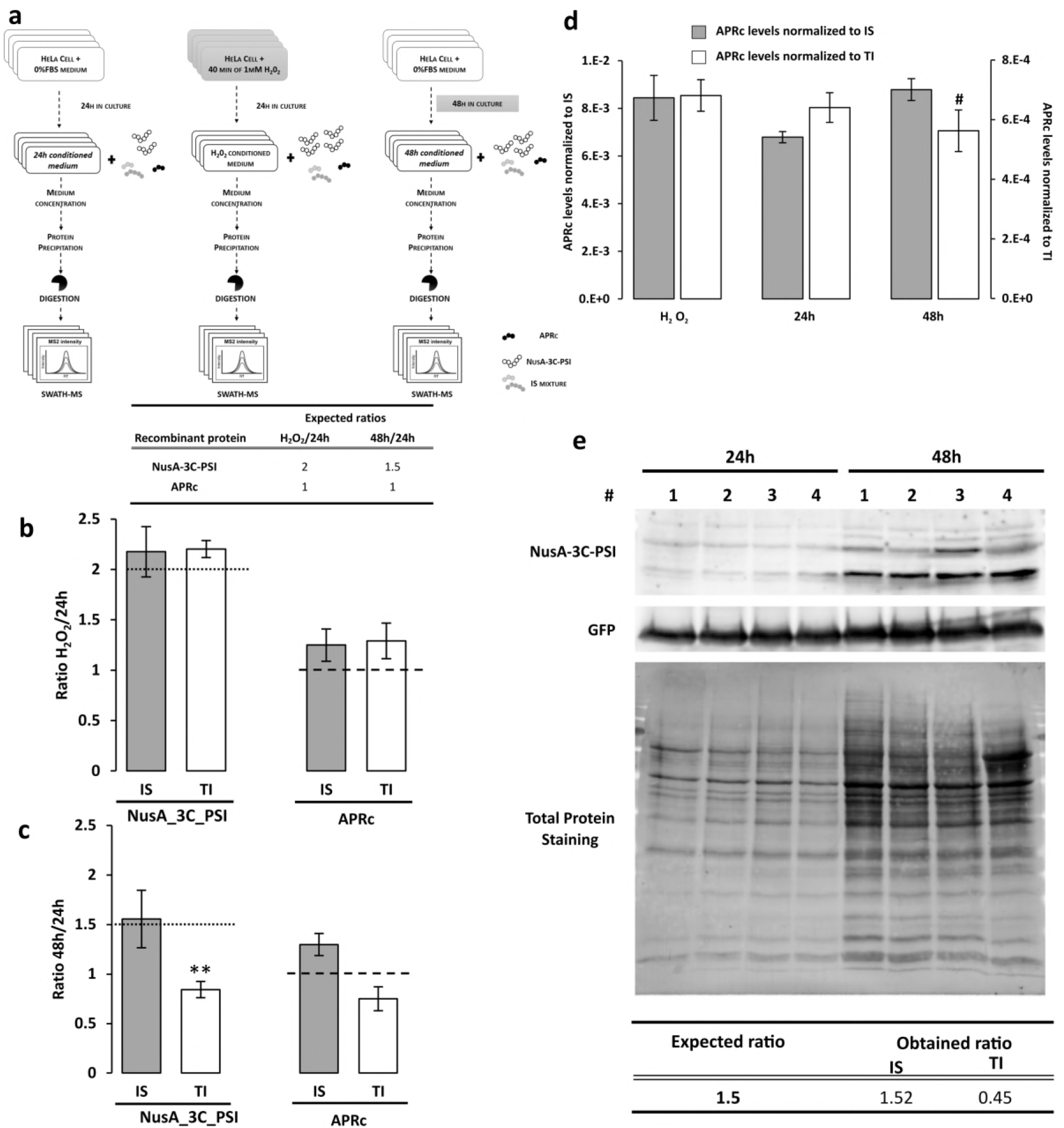
Evaluation of the normalization capacity of the proposed IS in a biologic context. **(a)** Experimental workflow. IS normalization capacity was confirmed by assessing the accuracy of the quantification of two recombinant proteins (APRc and NuSa-3C-PSI) spiked into different secretomes. The same amount of APRc was added to the three-conditioned media, while NuSa-3C-PSI was added in different proportions (bottom panel). The recombinant proteins and the IS were added immediately after the medium collection, and the entire samples were used for quantitative analysis. Secretomes with different complexities were obtained by performing experiments with and without an oxidant insult (H_2_O_2_) and two different conditioning periods (24h and 48h). The expected ratios for APRc (dashed lines) and NuSa-3C-PSI (dotted lines) are: 1 and 2, respectively, for the H_2_O_2_/24h comparison **(b)**; and 1 and 1.5, respectively, for 48h/24h comparison **(c)**. Data correspond to the mean ± SEM (n=4). \*\**ρ* <0.01, for significant differences to the respective expected ratio using a one-sample Student’s t-test. **(d)** Comparison of APRc levels among the three experimental conditions normalized using the IS (grey bars) or using the TI (white bars). Data represents the mean ± SEM (n=4). #*ρ*<0.05, for statistical significant differences for comparison among experimental conditions using Tukey HSD *post hoc* test. **(e)** Western blot validation. The NuSa-3C-PSI ratio between 48h and 24h was calculated after normalization using the IS or the total protein staining (TI). The expected and the mean ratios of four independent replicates are indicated in the panel below the representative blotting.

**Figure 4.**
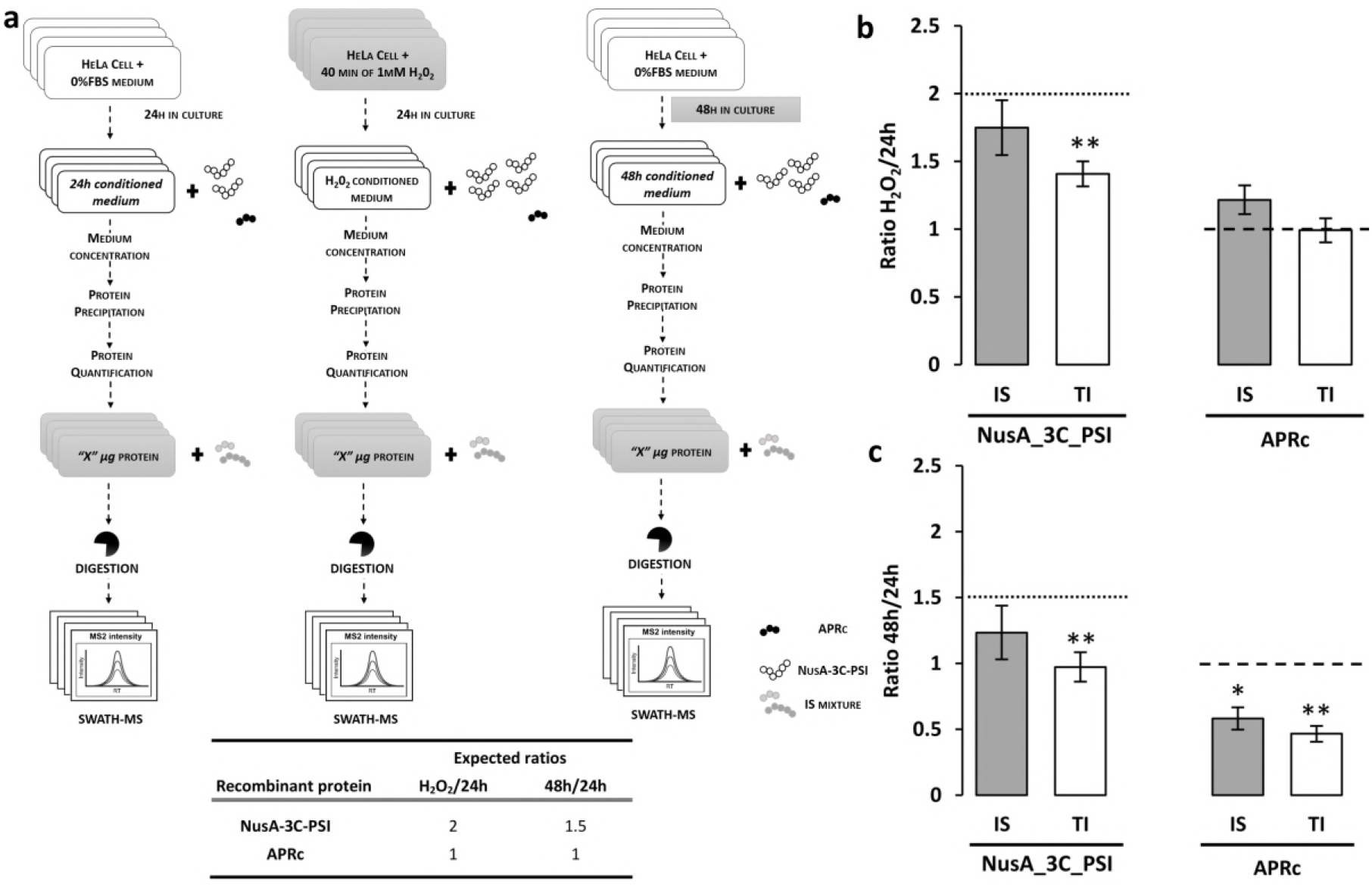
Analysis of protein quantification using the same amount of total protein content of samples with different degrees of complexity. **(a)** Experimental workflow used to confirm the impact of the analysis of the same amount of total protein for samples with large differences in sample composition, here exemplified by two different secretome analysis. The same experimental conditions as in Figure 3a were used to obtain secretomes with different complexities, and after the collection of the media, the two recombinant proteins (NuSa-3C-PSI and APRc) were spiked in the same proportions as those previously tested (and indicated in the bottom panel). Precipitated conditioned medium was quantified and the same amount of total protein was used in the quantitative analysis. The internal standard was only applied prior to the protein digestion step. The APRc and NuSa-3C-PSI levels were quantified using SWATH-MS/MS and normalized using the IS (grey bars) or TI (white bars), The fold change to the 24h condition was obtained and the expected ratios for APRc (dashed lines) and NuSa-3C-PSI (dotted lines) are: (i) 1 and 2, respectively, when compared the H_2_O_2_ condition with the control condition (H_2_O_2_/24h ratio) (b); and (ii) 1 and 1.5, respectively, for 48h/24h comparison (c). Data corresponds to the mean ± SEM of four independent experiments. \**ρ* <0.05 and \*\**ρ* <0.01, for significant differences to theoretical values (the respective expected ratio) using a one-sample Student’s t-test.

Based on the principle that the overall proteome content does not change [9], it is expected that the total intensity of the samples remain the same. Therefore, if any difference is observed this would reflect processing errors. This principle is particularly relevant for technical replicates as performed in these experiments; thus, as expected, the normalization to the total intensity (TI) was able to successfully reduce the intragroup variation observed in the raw data (Figure 2c), reducing from 75-100% coefficient of variation (CV) to values below 20% in the three conditions tested (considering all the 747 quantified proteins, including proteins quantified with a single peptide). The normalization using the proposed IS could reach a similar performance as the TI, revealing to be a good method to reduce intragroup variability.

A good internal standard should be able to reduce the variability between replicates, but also maintain the intergroup relations. To address this capacity, the ratios between the three experimental conditions with different complexities were evaluated. The fold change to the 30 μg condition was determined for all the proteins quantified with at least 3 peptides (proteins with highest confidence) and the obtained ratios were compared with the expected values of 1 and 3.33 for the TCA/30 μg and 100 μg/30 μg ratios, respectively (Figure 2d). In the case of the of the TCA/30 μg ratio, where only different samples processing methods were compared, the normalization for the total intensity leads to the expected results for at least 50% of the considered proteins (25^th^ to 75^th^ quartile). However, this method completely failed the 100 μg/30 μg ratio, to which the median fold change obtained was 1 instead of the expected 3.33. On the contrary, using the IS as normalization method it was possible to reach the two expected ratios. Thus, the proposed IS proves to be able to reduce the variability among replicates while maintaining the ratios between experimental conditions. Most importantly, the use of the IS not only proved to be beneficial in comparison to the total intensity but also when compared with a set of other 11 popular normalization methods (Figure 2b), which are able to largely decrease the variability between replicates (Supplementary Figure 5) but were not able to maintain the proportion between 30 and 100 μg of protein extract conditions (Figure 2e).

These results indicate that the use of IS may be particularly important in experiments that lead to large variations between conditions, including differences in sample complexity or differences in total protein content. The proteomics screenings of secretomes (combination of all the proteins secreted by cells under a particular condition, also known as conditioned medium) are a common example of experiments where the sample complexity may vary as a result of biological regulation. Therefore, secretome samples were used to confirm, in a biological context, the potential applicability of the IS for this type of challenging analyses, where a particular normalization method, such as total intensity, may fail.

Secretomes were obtained under different conditions (Figure 3a) that are usually tested in secretome studies, and are associated with differences in the total protein content: (i) the comparison between treated and untreated cells, herein using an oxidative stress insult caused by an acute stimulation with 1 mM of hydrogen peroxide; and (ii) time-course experiments, where the secretomes are collected after different cell culture periods. To evaluate if the normalization method leads to a correct quantification, two unrelated recombinant proteins (APRc, the retropepsin-like protease from *Rickettsia conorii* [17], and NusA-3C-PSI which is a fusion of NusA protein from *E. coli* with the saposin-like domain of a plant aspartic protease; Supplementary Figure 6) were added in known amounts to the secretomes immediately after medium collection to be used in targeted protein quantification in this analysis. These two recombinant proteins were added in low amounts (Supplementary Table 4), only enough to be quantified and to reflect the usual levels of the proteins analyzed. Additionally, while the same amount of APRc was added in the three conditions (24h or Control, 48h and H_2_O_2_ conditions), NusA-3C-PSI was added in distinct amounts in order to lead to two different ratios: 2 and 1.5-fold change for H_2_O_2_/24h and 48h/24h, respectively (Figure 3a). The internal standards were added immediately after the collection of the medium, and the entire sample was analyzed by mass spectrometry, maintaining the potential variation in the total protein content.

A prior proteomics characterization of these two recombinant proteins confirmed that they can be easily distinguished from the secreted proteins (Supplementary Figure 6 and Supplementary Tables 5 to 7) ensuring that a proper quantification can be achieved. The results obtained regarding the APRc and NusA-3C-PSI ratios confirmed previous results; while for H_2_O_2_/24h ratios (Figure 3b) both types of normalization lead to expected results (although some deviations are observed, these are not statistically significant), for the 48h/24h ratios (Figure 3c) only the normalization performed using the IS resulted in the expected values (indicated by the dotted and dashed lines for NusA-3C-PSI and APRc expected ratios, respectively). These two distinct results, particularly evident in the case of the NusA-3C-PSI, may be essentially associated with different degrees of sample complexity obtained within the three secretomes, as revealed by protein identification (Supplementary Figure 7) and most importantly by total protein quantification (Supplementary Table 8). A similar amount of total protein was obtained in the case of H_2_O_2_ and 24h allowing the use of the total intensity method. On the other hand, the total amount of protein in the 48h condition is twice the amount of that obtained at the 24h condition, impairing the use of methods based on the principle that the overall proteome content does not change, such as TI normalization method. Similar results to those obtained with TI were observed using other 11 methods proposed to be adequate for this analysis (Supplementary Figure 8), with a fold change near 1 being obtained for NusA-3C-PSI and APRc independently of the comparison performed (Supplementary Table 9), as well as for all the 772 proteins quantified in the three secretomes (Supplementary Figure 9). These results confirm the tendency of most of the normalization methods to buffer the intergroup differences, which in some cases can mask a biological result.

By being quantified with a lower number of peptides (only 3 peptides), the quantification of APRc is more prone to errors, which may justify the observed differences for the expected ratio of 1, independently of the normalization method used. To overcome that, and since the same amount of APRc was added to all samples, a statistical test was applied to compare the levels of APRc among the three secretomes (Figure 3d). These results, revealed that only the normalization using the IS confirms that no differences were observed between the three conditions. This further confirms that the normalization using the IS can be properly applied independently of sample complexity. On the other hand, with the TI normalization method, APRc was considered to be altered between the 48h and the H_2_O_2_ condition, substantiating the already observed indication that this normalization method is sensitive to substantial differences in sample complexity.

These results were validated by western blot (WB, Figure 3e and Supplementary Table 10) by determining the NusA-3C-PSI ratio among the 48h and 24h conditions. WB analysis revealed that the normalization to the IS (here only represented by GFP) was able to reach the expected ratio of 1.5, while the normalization to the total intensity (total protein staining) completely failed the expected ratio (calculated ratio of 0.45), confirming the results obtained in the MS analysis. Moreover, this result indicates that this IS strategy can be also used in other quantitative techniques such as western blot, when large differences in protein content are expected and no proper loading control is known for that particular application, such as in the case of secretome analysis.

Since the total protein content has a key impact for the selection of the sample normalization method, it was also evaluated if these differences may be taken into account in the analysis, by considering the entire sample, or the same volume of sample (as tested in the previous set of experiments Figure 3a) or if the differences in the total protein content may be diluted by analyzing the same amount of total protein (Figure 4a). To perform this evaluation, a similar set up was carried out for the addition of the recombinant proteins NusA-3C-PSI and APRc, but with the internal standard being added only after the protein quantification and adjustment of samples to the same amount of total protein. The two proteins were quantified and normalized using both IS and TI normalization methods, after which the different ratios were calculated (Figure 4b and c). In general, the use of the same amount of total protein, impairs the determination of the proper relations between the analyzed samples, independently of the normalization method: (1) in the case of the NusA-3C-PSI larger variations to the expected value (dotted line) were observed, in particular when the normalization to the total intensity is used; and (2) relatively to APRc, although no major differences were observed in the H_2_O_2_/24h ratio (consistent with the minor differences observed in total protein content between these two conditions, see Supplementary Table 8), the calculated ratio for the 48h/24h comparison completely failed the expected value of 1 (dashed line). These results indicate that the entire sample or the same volume of sample should be used to reflect the actual relations between samples, which is particularly important when large proteome variations are expected has a result of biological regulation.

Finally, to further evaluate the importance of performing a proper data normalization, the levels of the 769 endogenous proteins secreted upon different conditioning periods (24 and 48h) were evaluated using the two types of normalization (Figure 5). As already demonstrated for the two target proteins (APRc and NusA-3C-PSI, Figure 3) the normalization to the total intensity (Figure 5a) tends to eliminate the differences between the two samples (proteins ratios closer to 1, correspondent to the log2 of 0 in Figure5a), which resulted in the absence of proteins with a statistical meaningful alteration between the two-conditioned media. On the other hand, the normalization to the IS demonstrated to be more conservative of the actual differences between the two samples, with the volcano plot revealing a clear tendency for a large number of proteins being increased at the 48h conditioned medium (Figure 5b). This tendency is in accordance with the results obtained from the quantification of the protein total levels in each conditioned medium (Supplementary Table 8) which revealed an increase to the double of the average amount of protein of the 48h-conditioned medium compared to the CM obtained after 24h. Moreover, this analysis also revealed a group of 372 proteins presenting a statistically significant increase in the 48h condition, which is lost in the analysis using the normalization to the TI (Figure 5b and 5a, respectively). Those altered proteins are mainly associated with the extracellular milieu (Figure 5c) indicating a potential accumulation of secreted proteins/vesicles and a potential reorganization of the extracellular space [18].

**Figure 5.**
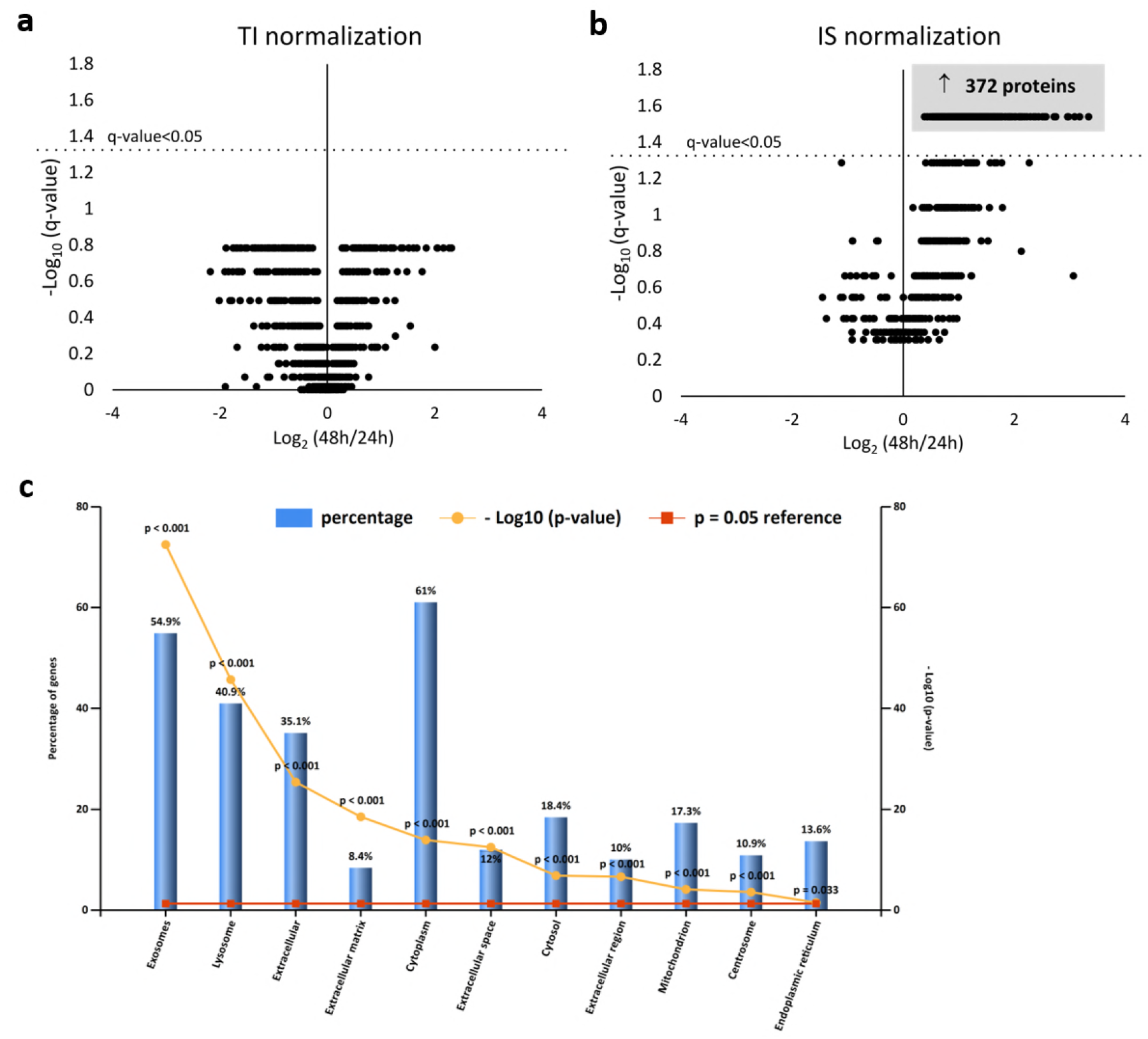
Analysis of the impact of data normalization for the identification of the proteins differentially secreted considering two different conditioning periods. Volcano plot reflecting the results from the statistical analysis of the 769 proteins quantified among two secretomes obtained after 24 or 48h of conditioning, using data normalized for the total intensity **(a)** or for the internal standard **(b)**. Statistical analysis was performed by Mann-Whitney and statistical significance was considered for q-values < 0.05 (i.e. using an adjusted p-value). **(c)** Cellular component analysis of the 372 proteins found statistically altered in (b) analysed by FunRich functional enrichment analysis (hypergeometric test p-value is indicated by the red line and Bonferroni corrected p-value is indicated in yellow with the corrected *p* < 0.05 meaning a significant enrichment).

## DISCUSSION

The introduction of the recombinant proteins GFP and MBP as IS for untargeted proteomics proved to be advantageous for the improvement of quantitative results obtained by SWATH-MS. The proposed internal standard strategy shows a similar intragroup normalization capacity when compared with the most common normalization methods, with the additional advantage of being able to maintain the actual proportions between conditions when large qualitative and quantitative differences in sample composition are observed, such as the ones induced by biological regulation or by enrichment approaches.

The herein proposed internal standard normalization approach proved to be a transversal method since: (i) it can be used for proteomics screenings of a wide range of species; (ii) it can be applied independently of the magnitude of the differences in the composition of the samples, being able to evaluate the dynamics of a targeted protein independently of total protein dynamics; and (iii) it can also be used with alternative techniques for validation of mass spectrometry assays, such as western blot. Moreover, the proposed IS method provides a more cost-effective and easier to implement alternative than the current stable-isotope labeling internal standards, being therefore an appealing strategy for large quantitative screenings, including biomarker discovery studies.

## METHODS

Methods and any associated references are available in the *online version of the paper* Note: Supplementary Information files are available in the *online version of the paper*.

## ACKNOWLEDGMENTS

This work was financed by the European Regional Development Fund (ERDF) through the COMPETE 2020 - Operational Programme for Competitiveness and Internationalisation and Portuguese national funds via FCT - Fundação para a Ciência e a Tecnologia, I.P., under projects: PTDC/NEU-NMC/0205/2012, POCI-01-0145-FEDER-007440 (ref.; UID/NEU/04539/2013), POCI-01-0145-FEDER-016428 (ref.: SAICTPAC/0010/2015), and POCI-01-0145-FEDER-016795 (ref.: PTDC/NEU-SCC/7051/2014); and by The National Mass Spectrometry Network (RNEM) under the contract ROTEIRO/0028/2013. S.I.A was supported by PhD fellowship SFRH/BD/81495/2011 co-financed by the European Social Fund (ESF) through the POCH - Programa Operacional do Capital Humano and national funds via FCT.

## AUTHOR CONTRIBUTION

S.I.A. performed the experiments and wrote the paper. I.S. and P.C. performed the recombinant protein production. All authors were involved in the design of the experiments and discussion of the results as well as in the revision of the manuscript.

## COMPETING FINANCIAL INTEREST

The authors have declared no conflict of interest.

## ONLINE METHODS

### Reagents

All reagents used in cell culture were cell culture-tested. The Dulbecco’s modified Eagle medium (DMEM) with Glutamax™ and low glucose (1 g/L), fetal bovine serum (FBS), trypsin 0.05% solution in phosphate buffered saline (PBS), Fungizone^®^ Antimycotic (amphotericin B), penicillin-streptomycin solution (Pen-Strep) solution, and Dulbecco’s phosphate buffered saline (DPBS) (10×) were obtained from Invitrogen. The hydrogen peroxide (H_2_O_2_) used in oxidative stress stimulation was obtained from Sigma-Aldrich.

Reagents used in the preparation of the diverse buffers for protein extraction, SDS-PAGE and immunoblotting have different sources. The tris(hydroxymethyl)aminomethane (Tris) was obtained from Calbiochem^®^ (Merck) and hydrochloric acid (HCl) was from JT Baker^®^. From BioRad: sodium dodecyl sulfate (SDS), dithiothreitol (DTT), acrilamide/bis-acrilamide solution [37.5:40 % (v/v)] and Precision Plus Protein™ - All blue standards. Glycine, ammonium persulfate, glycerol, bromophemol blue, Tween 20, and ECF reagent were from GE Healthcare. Sodium chloride (NaCl), magnesium chloride (MgCl_2_) and sodium dihydrogen phosphatase monohydrate were obtained from Merck, sodium-deoxycholate, potassium chloride (KCl), tetramethylethylenediamine (TEMED) and potassium phosphate dibasic trihydrate from Fluka (Sigma-Aldrich). IgePal, phenylmethylsulfonyl fluoride (PMSF), ethylenediaminetetraacetic acid (EDTA) and methanol were obtained from Sigma.

The reagents used in MS analysis were all high-quality chemical or reagents (ACS Reagent Chemicals & Lab Grades). Formic acid (FA) was obtained from AMRESCO and acetonitrile (ACN) from Fisher. Ortho phosphoric acid and ammonium sulfate were from Merck and ammonium bicarbonate from Fluka.

The Trypsin Modified Sequencing Grade used in protein digestion, the Complete Mini protease inhibitor mixture and Complete Mini phosphatase inhibitor mixture were obtained from Roche Diagnostics.

The source of the remaining reagents, antibodies and kits used in this work is referred throughout the text.

### Subproteome fractionation - membrane enrichment

The membrane enriched fractionation was obtained by ultracentrifugation performed as described in Anjo *el al*. [16, 19].

### Conditioned medium

HeLa cells were seeded at 12×10^3^ cell/cm^2^ in 55 cm^2^ plates (Corning) in a total of 4 plates per condition for mass spectrometry analysis or 1 plate for immunoblotting experiments. After 48 hours in culture (37 °C with 5% of CO_2_/95 % air and 95% humidity) the culture medium (DMEM medium with 10 % FBS) was discarded and cells were washed twice with warm PBS to remove the remaining FBS [20]. Then, the medium was changed for DMEM medium without FBS and cells were left in culture for 24 or 48 h (control conditions) or, cells were treated with 1 mM of H_2_O_2_ for 40 min to promote oxidative stress and then left in DMEM without FBS for 24 h.

After 24 h (ctrl and stress conditions) or 48 h of media conditioning, the medium was collected, centrifuged at 290 *xg*, for 5 min at 4 °C to remove cell debris, and then concentrated using cut-off filters of 5 kDa (Vivaspin20, Sartorius) [20]. The concentrated conditioned media were precipitated using Trichloroacetic acid (TCA) - Acetone as described in Manadas *et al*.,[21] and the protein pellets were re-suspended in 2× SDS Laemmli buffer.

### Recombinant proteins - Internal standard mixture and proteins used as targets in the normalization assays

#### Production of MBP and GFP Internal Standards

The DNA construct encoding for the fusion construct 6xHis-MBP-TEV-GFP (TEV - Tobacco Etch Virus protease cleavage site), cloned in pET28a vector was used to transform Escherichia coli BL21 star (DE3) strain. A single colony transformant was inoculated into 20 ml Luria Bertani (LB) medium containing 50 μg/ml and grown overnight at 37 °C. The culture was then transferred to 1L of fresh LB medium with 50 μg/ml kanamycin and allowed to grow at 37 °C until an OD600 of 0.7. Isopropyl-beta-D-thiogalactopyranoside (IPTG) was then added to a final concentration of 0.5 mM to induce protein expression at 37 °C. After 3 h, cells were harvested by centrifugation at 9000 xg for 15 min, resuspended in 20 mM Phosphate buffer containing 500 mM NaCl and 20 mM imidazole (buffer A) and cell lysis was obtained by 3 passages at 600-800 bar on an Avestin Emulsiflex C3 pressure homogeneizer, followed by ultracentrifugation (100 000 xg, 20 min) for extract clarification.

The fusion protein was purified on a AKTA chromatographic system (GE Healthcare Life Sciences), by immobilized metal ion affinity chromatography (IMAC) using a HisTrap HP, 5 ml column (GE Healthcare Life Sciences). The clarified cell extract was loaded into the column and buffer A was passed to remove unbound material until a stable horizontal baseline was obtained. Protein elution was performed by increasing the imidazole concentration stepwise (50, 100, 300, 500 mM imidazole), and fractions from each chromatography peak were analyzed by SDS-PAGE. The fractions containing the fusion protein were pooled and buffer was exchanged to 25 mM Tris-HCl pH 8.0, 150 mM NaCl using a 5 ml HiTrap Desalting column (GE Healthcare Life Sciences).

Protein was digested by incubation with TEV protease in a 1:50 ratio (w/w) at 4 °C, overnight. The digested protein was passed through a 5 ml MBPTrap HP column (GE Healthcare Life Sciences) to remove MBP, which was eluted by passing 10 mM maltose in PBS buffer, yielding purified MBP. The MBPTrap flow-through containing the GFP and TEV protease was passed through a negative IMAC to remove the TEV protease which had a N-terminus hexahistidine tag, yielding purified GFP.

After purification, protein purity was assessed by SDS-PAGE and the internal standard was prepared by mixing the proteins in a 1:1 ratio (w/w). Protein was quantified using the 2-D Quant kit (GE Healthcare) according to the manufacturer’s instructions, and the samples were stored at −20 °C until further use.

#### Production of NusA-3C-PSI and APRc

The cDNA of plant specific insert (PSI) domain of the plant aspartic protease cardosin A [22] was cloned into pCoofy 16 vector [23], coding for a fusion protein contain an N-terminal decahistidine tag, followed by NusA protein, a human rhinovirus 3C protease cleavage site and the PSI domain at the C-terminus.

Protein expression and extract preparation was performed as for the Internal Standard. The fusion protein was purified by IMAC using a HisTrap HP, 5 ml column (GE Healthcare Life Sciences). Protein elution was performed by increasing the imidazole concentration stepwise (50, 100, 300, 500 mM imidazole), and fractions from each chromatography peak were analyzed by SDS-PAGE. The fractions from the peak with purified protein were combined and the buffer exchanged to PBS using a 5 ml HiTrap Desalting column (GE Healthcare Life Sciences).

APRc was produced as indicated in Cruz *et al*., [17].

Recombinant proteins were quantified using the 2-D Quant kit (GE Healthcare) according to the manufacturer’s instructions. All samples were stored at −20 °C until further use.

### Analysis of the reproducibility of the Internal standard

For this analysis, the samples were subjected to liquid digestion as described in Anjo *et al*., [16]. Several conditions were performed to evaluate the impact of the each of the steps usually performed in a mass spectrometry approach (see Figure 1a).

For reproducibility assays at MS and sample clean-up level, 400 μg of the membrane-enriched sample and 50 μg of internal standard (IS) mixture (MBP and GFP) were subjected to liquid digestion in separate, and their mixture was performed at peptide level prior to LC-MS/MS analysis or prior to C18 cleanup in a ratio of 1 μg of IS peptides per 30 μg of extract peptides. For MS analysis, the IS peptides alone and a single mixture of IS and extract peptides were prepared and subjected to four SWATH-MS acquisitions. For the C18 cleanup test, a single mixture of 120 μg of extract peptides and 4 μg of IS peptides was prepared, and then processed in four independent micropipette stage tips. In a similar way, four batches of 10 μg of the same IS peptide mixture were processed in independent micropipette stage tips.

To evaluate the impact of digestion, sample complexity, and sample processing on IS reproducibility, four replicate digestions were performed for the following samples: (i) 10 μg of IS alone, (ii) a mixture of 30 μg of membrane enriched extract spiked with 1 μg of IS, (iii) a mixture of 100 μg of membrane enriched extract spiked with 1 μg of IS, and (iv) a mixture of 30 μg of membrane enriched extract spiked with 1 μg of IS subjected to TCA/acetone precipitation.

All samples were resuspended in LC-MS/MS mobile phase (2% ACN and 0.1% FA) spiked with iRTs peptides (Biognosys AG) to a final concentration of 1 μg IS/30 μL of sample. Samples were analyzed on an AB Sciex 5600 TripleTOF in two modes: information-dependent acquisition (IDA) for protein identification and library generation, and SWATH acquisition for quantitative analysis (see detailed information in the Supplementary Files - SWATH-MS method 1).

Information dependent acquisition (IDA) experiments were performed for a representative sample of the membrane enriched extract spiked with the IS mixture (MBP and GFP proteins), and for the IS alone. A single injection of 10 μL of each sample was set for quantitative analysis by acquisition in SWATH mode. The SWATH setup was essentially as in Gillet *et al*. [24], with the same chromatographic conditions used in the IDA experiments.

Protein identification of each sample was obtained using ProteinPilot™ software (v4.5, AB Sciex^®^) with the following search parameters: search against the entire SwissProt database (released at February 2014) and iRT peptides sequences; trypsin digestion; and MMTS as cysteine alkylating reagent. The protein identification files were then used to create a specific library of precursor masses and fragment ions of each sample to be used for subsequent SWATH processing. An independent False Discovery Rate (FDR) analysis, using the target-decoy approach provided by Protein Pilot™, was used to assess the quality of identifications. Positive identifications were considered when identified proteins and peptides reached a 5 % local FDR [25, 26].

SWATH data was processed in the SWATH™ processing plug-in for PeakView™ (v2.0.01, AB SCIEX). Briefly, the chromatographic profiles of the peptides presented in the library were extracted from the SWATH-MS data (in an extracted-ion chromatogram (XIC) window of 5 min) for up to 5 target fragment ions of up to 15 peptides per protein. Confident identification was considered for proteins with at least one peptide with a FDR below 1%. Additionally, the retention time was adjusted to each sample using the iRT peptides. Protein levels were estimated by summing all the transitions from all the peptides of a particular protein. [16]

### Evaluation of the normalization capacity of the internal standard (IS)

The recombinant proteins NuSa-3C-PSI and APRc were subjected to liquid digestion (as described in [16]) for their characterization by LC-MS/MS prior to the study of IS normalization capacity. Briefly, 10 μg of each protein were digested with 0.2 μg trypsin (0.1 μg/μL in 0.5 TEAB) overnight at 37 °C.

The conditioned medium (CM) samples were analyzed in two different modes for a total of four biological replicates each: (1) using the same volume between samples or (2) the same amount of total protein per sample. In both analyses the recombinant proteins NuSa-3C-PSI and APRc were added to the medium prior to medium concentration according to the Supplementary Table 4, while the IS mixture was added prior to concentration in the case of the “same volume” analysis (4 μg of IS per sample), and before protein digestion in the case of the “same amount” analysis (2 μg of IS per sample) (Figure 3a and 4a, respectively).

In the “same volume” analysis, after concentration and precipitation of the protein the entire sample dissolved in 2x SDS Sample buffer were used for in-gel digestion. On the other hand, in the case of the analysis using the “same amount of total protein amount”, after dissolution of the protein pellets, samples were quantified using the 2-D Quant kit (GE Healthcare) according to the manufacturer’s instructions, and 30 μg of total protein were used for protein digestion. Additionally, one plate of each biological replicate was combined to create a pooled sample for each CM condition. These pooled samples were spiked with the recombinant proteins and digested using the same condition of the individual replicates, and were used for IDA experiments to build a specific protein library to be used in SWATH-MS analysis.

Protein digestion was performed using the Short-GeLC approach [16] with slight modifications. Briefly, each lane was divided into 3 fractions of similar size, and processed individually. After in-gel digestion the formed peptides from each fraction were pooled together into a single sample in the case of the samples for SWATH analysis, or maintained separate in the pooled samples for IDA experiments. CM samples from the “same volume” experiments were resuspended in 13 μL of mobile phase and the CM samples from the “same amount” experiments in 24 μL.

Samples were analyzed by SWATH-MS mode as described above but with some modifications (see detailed information in the Supplementary Files - SWATH-MS method 2). IDA experiments were performed for each fraction of a representative pooled sample of each CM condition spiked with the recombinant proteins used in the experiment (GFP and MBP (IS), NuSa-3C-PSI and APRc). NuSa-3C-PSI and APRc individual samples were also subjected to IDA experiments for protein characterization.

In these experiments, the SWATH-MS setup was designed specifically for the samples to be analyzed (Supplementary Table 11), to that a pool of all samples was analyzed in IDA mode in order to be used to adapt the SWATH windows to the complexity of the set of samples to be analyzed.

Protein identification was obtained using ProteinPilot™ software (v5.0, AB Sciex^®^). NuSa-3C-PSI and APRc were analyzed using the following search parameters: search against the entire SwissProt database (released at October 2014) or against a sequence specific database; trypsin digestion; and MMTS as cysteine alkylating reagent. Conditioned medium samples were combined in a single search against a database composed by the *Homo sapiens* database from SwissProt (released at August 2014) and the sequences of all the recombinant proteins spiked (MBP and GFP (IS), NuSa-3C-PSI and APRc), using trypsin and acrylamide as alkylating agent, and a special focus option for gel-based approaches. Positive identifications were considered when identified proteins and peptides reached a 5 % local FDR [25, 26]. The identification file of the CM samples was used as the specific library of precursor masses and fragment ions for subsequent SWATH processing.

SWATH data was processed in the SWATH™ processing plug-in for PeakView™ (v2.0.01, AB SCIEX). Briefly, the chromatographic profiles of the peptides presented in the library were extracted from the SWATH-MS data (in an extracted-ion chromatogram (XIC) window of 4 min) for up to 5 target fragment ions of up to 15 peptides per protein. Confident identification was considered for proteins with at least one peptide with a FDR below 1%. Additionally, the retention time was adjusted to each sample using the IS peptides. Protein levels were estimated by summing all the transitions from all the peptides of a particular protein. [16]

The mass spectrometry proteomics data have been deposited to the ProteomeXchange Consortium via the PRIDE [27] partner repository with the dataset identifier PXD009068”

### Western blotting validation

Twenty-four and 48 h conditioned media were spiked with 0.25 μg of the IS solution (MBP and GFP recombinant proteins) and different amount of NuSa-3C-PSI and APRc according with the Supplementary Table 12 prior to culture medium concentration.

The samples were denatured by boiling at 95°C for 5 min, and the entire volume was loaded per lane and separated on 12.5% SDS-polyacrylamide gels using a mini-PROTEAN^®^ Tetra Electrophoresis System (Bio-Rad). Proteins were transferred to low fluorescence polyvinylidene fluoride (PVDF) membranes (TBT RTA TRANSFER KIT, Bio-Rad) using a Trans-Blot Turbo Transfer System (BioRad) during 45 min at a constant voltage of 25 V (with the current limited to 1 A). Following transfer, the membranes were blocked for 1 hour at room temperature (RT) with 5% (w/v) skimmed milk powder in PBS-Tween 20 (PBS-T) [0.1% (v/v)]. The membrane was incubated sequentially with anti-histidine, anti-APRc and anti-GFP (Supplementary Table 13) overnight at 4°C followed by 1 h at RT, prepared in the blocking solution. Primary antibodies were removed, and membranes were extensively washed with PBS-T (3 times, 15 min under agitation each time). Blots were then incubated for 1 h at RT with the respective secondary antibodies conjugated with alkaline phosphatase (Supplementary Table 14) in 5% (w/v) skimmed milk powder dissolved in PBS-T followed by extensive washes as above. The membrane was firstly incubated with the histidine specific antibody (to recognition of NuSa-3C-PSI protein) followed by re-probing of the membrane with antibody against APRc, and finally with a GFP specific antibody.

Protein-immunoreactive bands were developed using the “Enhanced Chemifluorescence (ECF) detection system” (GE Healthcare) and visualized in a Molecular Imager FX System (Bio-Rad). For determination of the total intensity of the sample loaded, the membrane was further stained using the LavaPurple Total Protein Stain kit (Gel Company) according with the manufacturer’s instruction for staining of PVDF membranes. Briefly, the membrane was re-wetted in methanol and rinsed in water prior to the protein staining, and then incubated in the staining solution composed by 125 μL of LavaPurple reagent in 50 mL of staining buffer (provided in the kit) for 30 min at RT. After incubation, the membrane was transfer to the fixation/acidification solution (provided in the kit) for 5 min at RT and rinsed three times in 100 % ethanol for 2-3 min at RT. After staining, the membrane was dried, and the signal was visualized in a Molecular Imager FX System (Bio-Rad) using the Deep Purple Filter.

The adjusted volumes (total intensities in a given area with local background subtraction) for each band and the total intensity of each lane were obtained using the Image Lab software (version 5.1, Bio-Rad).

### Data analysis

Recombinant proteins, both the internal standard mixture of MBP and GFP, NUSA-3C-PSI and APRc, were characterized by a series of bioinformatics and proteomics analyses in order to evaluate the similarity of these proteins with other proteins present in the reviewed database from UniProt. To evaluate sequence similarity, sequence alignment analysis was performed by using the Basic Local Alignment Search Tool (BLAST) from UniProt, and the obtained data was evaluated by the analysis of the calculated expectation values (E-values) and percentage of identity. On the other hand, protein identification was used to detect the proteins related with the recombinant proteins used in this work. These related proteins, which correspond to proteins that can be identified by the same set of peptide(s) (shared peptides), are grouped and reported as Proteins Groups (N) in Protein Pilot software and were further confirmed by the analysis of the Unused value, which corresponds to the ProScore contribution of the unique peptides, and determines if there is enough evidence to report a given protein. Protein identification was considered for all the Protein Groups that met the 5 % local FDR.

The consistency of the quantification (robustness) of the proposed IS mixture and the impact of sample complexity were determined by the analysis of the coefficient of variation (CV) of the (i) technical replicates, (ii) between the “-/+ extract” conditions from the same experiment, and ultimately (iii) between all the experimental conditions. For CV analysis, the IS values were normalized by the total intensity of the elements common to all samples [the internal standard and the iRT peptides (Biognosys) spiked prior to LC-MS/MS analysis] to reduce the expected technical variation associated with, for instance, difference in the sample or injection volume.

The normalization capacity was evaluated by (i) the capacity to reduce the intragroup variability through the analysis of the coefficient of variation between replicates, and (ii) by the capacity to reach the expected ratios. The normalization using the IS mixture was compared with a set of different well-established methods [12], with a special focus for the total intensity (TI) method. The quantification of the IS was obtained by summing the MBP and GFP levels, and the sample intensity was calculated by summing the levels of all the quantified proteins. The remaining normalizations were performed using the web tool “Normalyzer” (version 1.1.1), available at http://normalyzer.immunoprot.lth.se/ [12] that performs the evaluation using 12 different normalization methods, widely used in untargeted proteomics, and three different strategies: i) pooled coefficient of variation (PCV); ii) pooled median absolute deviation (PMAD); and iii) pooled estimate of variance (PEV). The normalization methods considered are: Log2, <u>LO</u>cally <u>WE</u>ighted <u>S</u>catter-plot <u>S</u>moothing (LOWESS or LOESS), Robust Linear Regression (RLR), Variance Stabilizing Normalization (VSN), total intensity (TI), median intensity (MedI), average intensity (AI), NormFinder (NF) and quantile. In the case of VSN, RLR and LOESS, the normalization can be achieved at global (G) or at local/replicate (R) level. Global normalization is performed across all samples without considering any replicate grouping, while local normalization is only performed within the replicates from the same group.

Box plots and Violin plots were used to represent the distribution of large sets of data. In the box plots, the limits of the box indicate the 25^th^ and 75^th^ percentiles, and Tukey-whiskers extend 1.5 times the interquartile range from the 25^th^ and 75^th^ percentiles, after which the outliers’ values are indicated. In the Violin plots, the median of the data is indicated (indicated by a white circle), the 25^th^ and 75^th^ percentiles of the data limited by the box, the non-outliers range of values indicated by the Tukey-whiskers extended 1.5 times the interquartile range from the 25^th^ and 75^th^ percentiles, and finally an estimation of the data distribution calculated by the Kernel density estimation which extend to extreme values represented by a polygon.

Protein quantification analysis of the recombinant proteins NuSa-3C-PSI and APRc using the two different normalization methods (IS and TI) was verified by statistical analysis using the one-sample Student’s t-test against a theoretical value correspondent to the expected ratio, and the ANOVA (ANalysis Of Variance) analysis followed by the Tukey HSD *post hoc* test for comparison of APRc levels among the experimental conditions. Significant differences were considered for p-values below a significance level of 5 %. Data presented as mean ± standard error of the mean (S.E.M.). Every experimental condition was tested in four sets of independent experiments. Parametric assumptions (data normality and homogeneity of variance) were tested using Shapiro-Wilk Test and Levene’s Test, respectively. Statistical analysis was performed using IBM^®^ SPSS^®^ Statistics Version 22.

Differential secretome analysis was performed using the results from the analysis of the same volume of sample from the experiment correspondent to the two conditioning periods (24 and 48h). Protein total levels were normalized for the total intensity or the internal standard. The proteins altered between the two conditions were identified by the Mann-Whitney test performed in InfernoRDN (version 1.1.5581.33355) [28] using the normalized values, and statistical significant differences were considered for q-values < 0.05.

Functional enrichment analysis was performed with FunRich (version 3.1.3) using FunRich human database, and statistically analyzed with hypergeometric test using FunRich human genome database as the background [29]. A Bonferroni corrected p-value < 0.05 indicates a sub-group of proteins that is significantly enriched in the sample against the background database.

